# Differential trafficking of albumin and IgG facilitated by the neonatal Fc receptor in podocytes in vitro and in vivo

**DOI:** 10.1101/494823

**Authors:** James Dylewski, Evgenia Dobrinskikh, Linda Lewis, Pantipa Tonsawan, Parmjit S. Jat, Judith Blaine

## Abstract

Proteinuria is strongly associated with kidney disease progression but the mechanisms underlying podocyte handling of serum proteins such as albumin and IgG remain to be elucidated. We have previously shown that albumin and IgG are transcytosed by podocytes in vitro. In other epithelial cells, the neonatal Fc receptor (FcRn) is required to salvage albumin and IgG from the degradative pathway thereby allowing these proteins to be transcytosed or recycled. Here we directly examine the role of FcRn in albumin and IgG trafficking in podocytes by studying handling of these proteins in FcRn knockout (KO) podocytes in vitro and in a podocyte-specific FcRn knockout mice in vivo. In vitro, we find that knockout of FcRn leads to IgG accumulation in podocytes but does not alter albumin trafficking. Similarly, in vivo, podocyte-specific knockout of FcRn does not result in albumin accumulation in podocytes in vivo as measured by mean albumin fluorescence intensity whereas these mice demonstrate significant intraglomerular accumulation of IgG over time. In addition we find that podocyte-specific FcRn KO mice demonstrate mesangial expansion as they age and activation of mesangial cells as demonstrated by increased expression of α-smooth muscle actin. Taken together, these results suggest that trafficking pathways for albumin and IgG differ in podocytes and that sustained disruption of trafficking of plasma proteins alters glomerular structure.

## Introduction

Proteinuria is an independent marker of kidney disease progression and is widely used clinically as a biomarker of kidney dysfunction (1, 2). Proteinuria is both a consequence of kidney damage and damages the glomerulus and tubules directly by increasing production of pro-inflammatory cytokines and promoting fibrosis (1, 3-5). Both the glomerulus and the proximal tubules are involved in the renal handling of serum proteins but the molecular mechanisms remain to be fully elucidated. The primary barrier to filtration of large plasma proteins into the urine is the glomerular filtration barrier (GFB) which consists of three layers – a fenestrated endothelium, the glomerular basement membrane and the podocyte (6). The podocyte is a specialized epithelial cell containing a large cell body and multiple processes which ramify to form smaller processes. Paracellular passage of large serum proteins is prevented by the slit diaphragm which extends between the foot processes of neighboring podocytes and precludes filtration of proteins ∼ 70 kDa or larger. The precise amount of albumin filtered through the GFB is a contested topic (7, 8). By even the most conservative estimates, ∼ 4 g albumin a day transit the GFB (9). The amount of IgG that traverses the glomerular filtration barrier is unknown.

Podocytes have been shown to take up albumin in vitro and in vivo (4, 10, 11). Using in vitro assays, we have previously shown that podocytes endocytose albumin and that most is transcytosed, with a smaller amount sent to the lysosome for degradation (12). These findings have been confirmed in vivo using intravital multiphoton microscopy in rats (11). In other epithelial cells, including those in the renal proximal tubule, the neonatal Fc receptor (FcRn) is required to prevent albumin and IgG from entering the degradative pathway, thereby allowing albumin to be recycled or transcytosed (13-16). FcRn, has homology to major histocompatibility complex class I and binds albumin and IgG at pH 6-6.5 but has minimal affinity for these proteins at neutral pH (17). Within the adult kidney, FcRn is expressed in podocytes, vascular endothelial cells and the proximal tubule (18). The physiologic role of FcRn in albumin trafficking in podocytes is unknown.

Akilesh et al. demonstrated that the neonatal Fc receptor is required to prevent the intraglomerular accumulation of IgG in mice (19). These studies were performed in global FcRn knockout (KO) mice which manifest hypoalbuminemia and hypogammaglobulinemia. Plasma levels of albumin and IgG are 50% (16) and 80-90% (20) lower respectively in global FcRn KO compared to wild type (WT) mice. Thus podocytes in the global KO are exposed to significantly less albumin and IgG than WT mice which might alter trafficking pathways.

Here we use in vitro assays and podocyte-specific FcRn knockout mice to directly examine the role of FcRn in albumin and IgG trafficking in podocytes. Creation of podocyte-specific FcRn KO mice allowed for the examination of intraglomerular trafficking of albumin and IgG in mice that have normal serum levels of these proteins, permitting direct assessment of FcRn mediated trafficking of albumin and IgG in podocytes.

## Results

### Establishing WT and FcRn KO podocyte cell lines

In order to directly investigate the role of FcRn in albumin and IgG trafficking in podocytes, we isolated podocytes from WT and global FcRn KO mice and created conditionally immortalized cell lines by transforming primary podocytes with the thermosensitive SV40 T antigen (21). Differentiated WT and FcRn KO podocytes expressed the podocyte markers podocin and Wilms tumor 1 (WT1) (Figure 1A) and FcRn KO podocytes had minimal FcRn mRNA as assessed by qPCR (Figure 1B). FcRn knockout did not impair albumin or IgG uptake in podocytes (Figure 1C).

**Figure 1.**
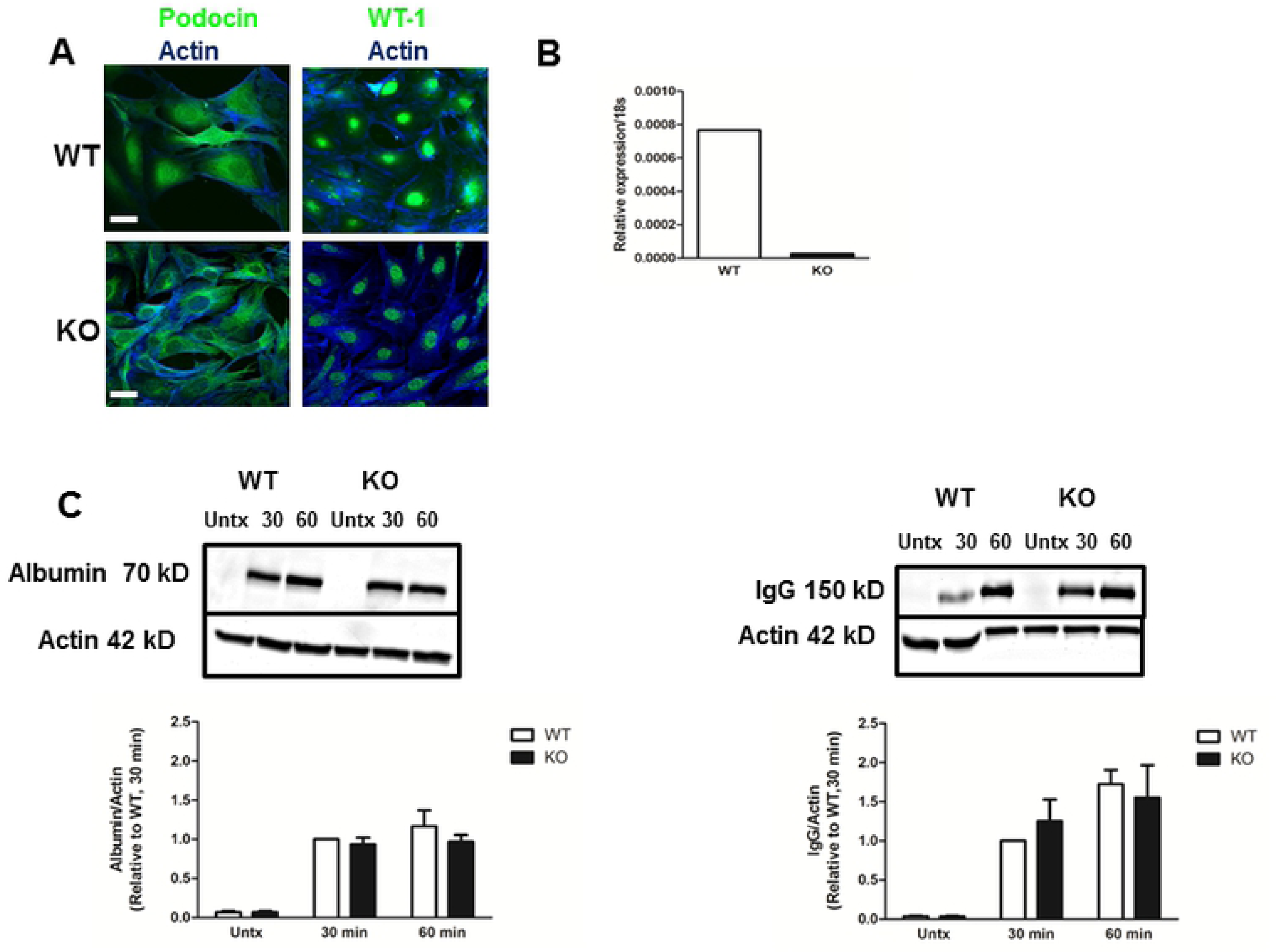
*Characterization of FcRn knockout podocytes*. *A*, Conditionally immortalized wild type (WT) and FcRn knockout (KO) podocytes express the podocyte markers podocin and Wilms Tumor 1 (WT1). Scale bar 20 um. *B*, FcRn KO podocytes have minimal expression of FcRn mRNA. *C*, There was no significant differences in uptake of FITC-albumin or IgG in wild type versus FcRn KO podocytes. n = 3 experiments.

### FcRn KO impairs IgG trafficking but not albumin trafficking in vitro

To examine IgG and albumin trafficking in vitro, differentiated WT and FcRn KO podocytes were loaded with albumin or IgG at 4 °C (a temperature that allows surface binding of albumin or IgG but inhibits endocytosis) or 37° C (permits endocytosis), washed well after loading and harvested immediately or at the times indicated in Figure 2. Leupeptin, a lysosomal enzyme inhibitor, was also used to examine the effects of blocking lysosomal degradation on albumin and IgG trafficking in WT and FcRn KO podocytes. In WT podocytes loaded with IgG at 37°C, there was a decrease in the amount of IgG remaining in the cells after 30 minutes incubation in Ringer solution (Figure 2A). Inhibition of lysosomal degradation with leupeptin did not significantly increase the amount of IgG remaining in WT podocytes after 30 minutes incubation in Ringer solution, suggesting that monomeric IgG is not trafficked to the lysosome. In contrast to WT podocytes, the amount of IgG remaining intracellularly in FcRn KO podocytes 30 minutes after loading with IgG was not significantly decreased, suggesting impairment in IgG transcytosis in the KO (Figure 2A).

**Figure 2.**
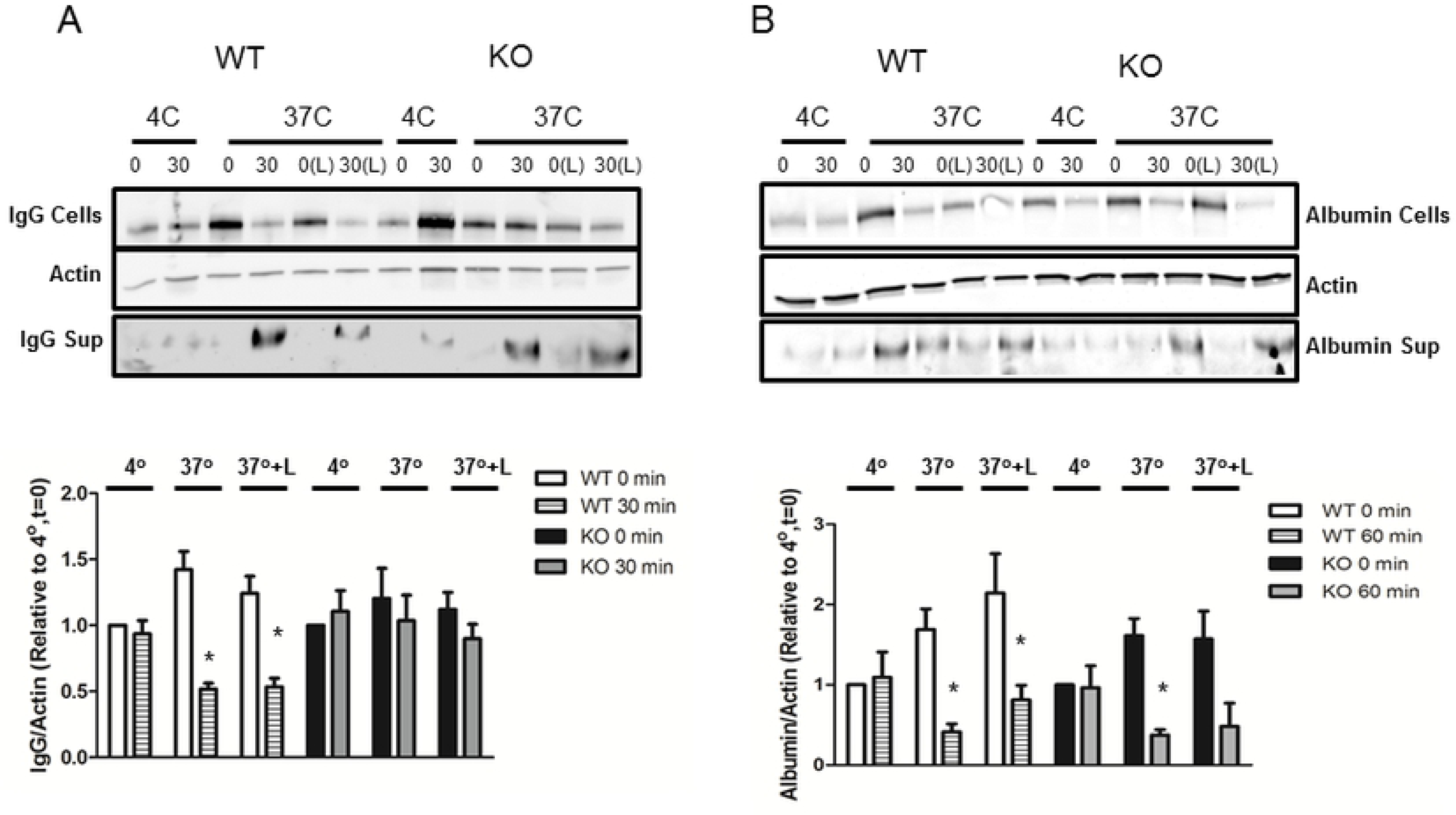
*FcRn KO impaired albumin but not IgG transcytosis in podocytes in vitro*. *A*, Cultured podocytes from FcRn KO mice demonstrate impaired IgG transcytosis. There is significantly less monomeric IgG in the cellular fraction in WT podocytes 30 min after loading with IgG, whereas the amount of monomeric IgG in the cellular fraction in KO podocytes is comparable to that at t = 0. Inhibition of lysosomal degradation does not alter the amount of monomeric IgG present in the cellular fraction suggesting that IgG is not sent to the lysosome. *, p = 0.0017 compared to the same condition at t = 0; n = 10 experiments. Time (0, 30) is in minutes; L = leupeptin. *B*, In contrast, FcRn KO has no effect on albumin transcytosis in cultured podocytes. There is significantly less albumin in the cellular fraction in both WT and KO podocytes 60 minutes after loading with albumin. *, p < 0.0001 compared to the same condition at t = 0; n = 8 experiments. Time (0, 60) is in minutes; L = leupeptin.

Albumin trafficking in WT and FcRn KO podocytes was examined by loading podocytes with FITC-labeled albumin, washing the podocytes very well and then assessing how much FITC-albumin remained in the podocytes immediately after washing or after 60 minutes in Ringer solution ± leupeptin to inhibit lysosomal degradation. Podocytes were assessed after 60 min as initial studies demonstrated that albumin is trafficked more slowly than IgG. As shown in Figure 2B, in both WT and FcRn KO podocytes loaded with FITC-albumin at 37 °C there was a significant decrease in the amount of albumin remaining in the cells 60 minutes after incubation in Ringer solution. Thus, knockout of FcRn did not impair albumin trafficking in podocytes in vitro. In both WT podocytes, there was a trend towards increased albumin accumulation in leupeptin treated cells at t= 60 minutes, 37° C but this was not significant

### Podocyte-specific FcRn KO mice

In order to determine whether the differential trafficking of albumin and IgG in podocytes lacking FcRn occurred in vivo, we generated podocyte-specific FcRn KO (FcRn fl/fl;Podocin-Cre/+) mice by crossing podocin-Cre mice with FcRn floxed mice (Figure 3). FcRn floxed mice lacking the Cre transgene (FcRn fl/fl;+/+) served as littermate controls. FcRn fl/fl;Podocin-Cre/+ mice had similar serum levels of albumin and IgG compared to controls (Figure 3A). There was no difference in the urinary albumin to creatinine ratio in podocyte-specific FcRn KO mice versus control at 3, 6 and 12 months (Figure 3B).

**Figure 3.**
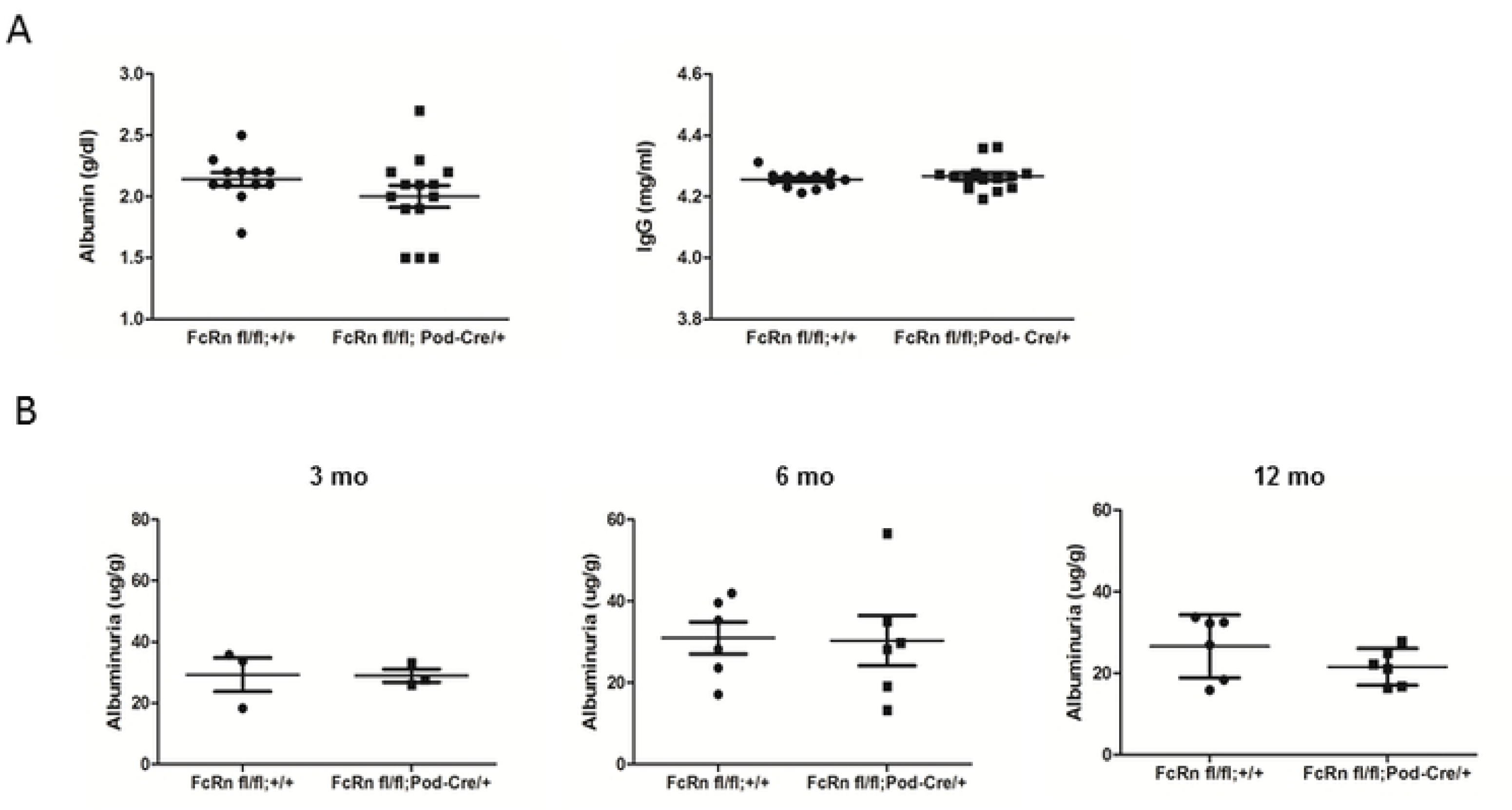
*Functional parameters in podocyte-specific FcRn knock-out mice*. *A*, There were no differences in serum albumin or IgG levels between podocyte-specific FcRn KO (FcRn fl/fl; Pod-Cre/+) and control (FcRn fl/fl;+/+) mice (n= 14 KO and n = 12 control mice). *B*, Podocyte-specific FcRn KO and control mice had minimal albuminuria at 3 months and no significant increase in albuminuria with age (n = 3 control and 3 KO mice at 3 months, 6 KO and 6 control mice at 6 months and 6 KO and 6 control mice at 12 months).

### Podocyte-specific KO of FcRn results in IgG accumulation in the glomerulus

We used immunofluorescence staining to examine whether lack of FcRn in podocytes results in albumin or IgG accumulation in the glomerulus as mice age. There was no significant difference in IgG accumulation in the glomeruli of podocyte-specific FcRn KO versus control mice at 3 months of age (mean IgG fluorescence/glomerular area 3.9 ± 0.5 vs 5.1 ± 0.4, p = NS; Figure 4A). By 6 months of age, podocyte-specific FcRn KO mice had a significant increase in glomerular IgG compared to controls (5.7 ± 0.3 vs 4.1 ± 0.2, p < 0.05). At 12 months of age there was a statistically significant increase in the amount of IgG present in the glomerulus in podocyte-specific FcRn KO compared to controls (9.4 ± 0.5 vs 5.6 ± 0.5, p < 0.0001).

**Figure 4.**
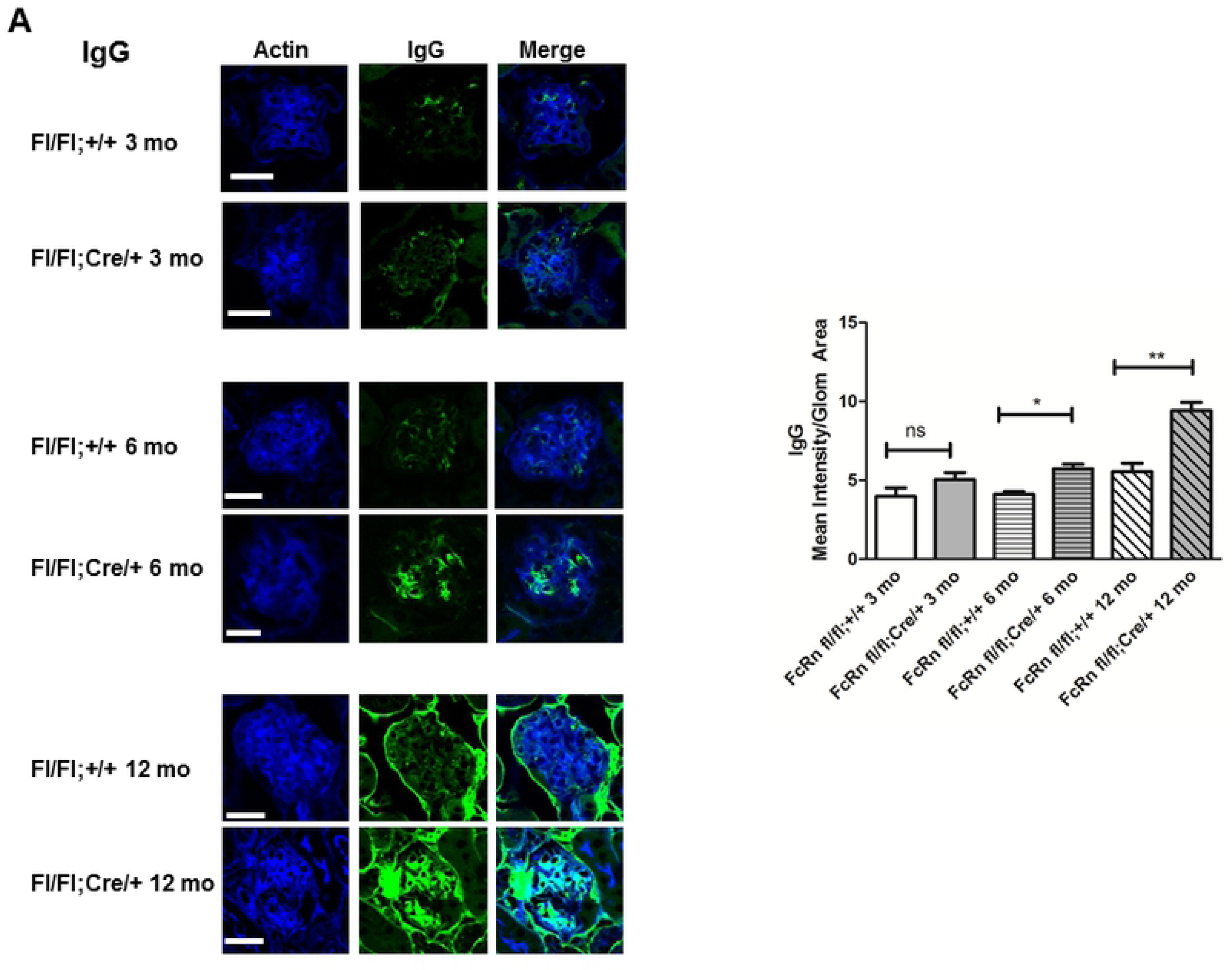
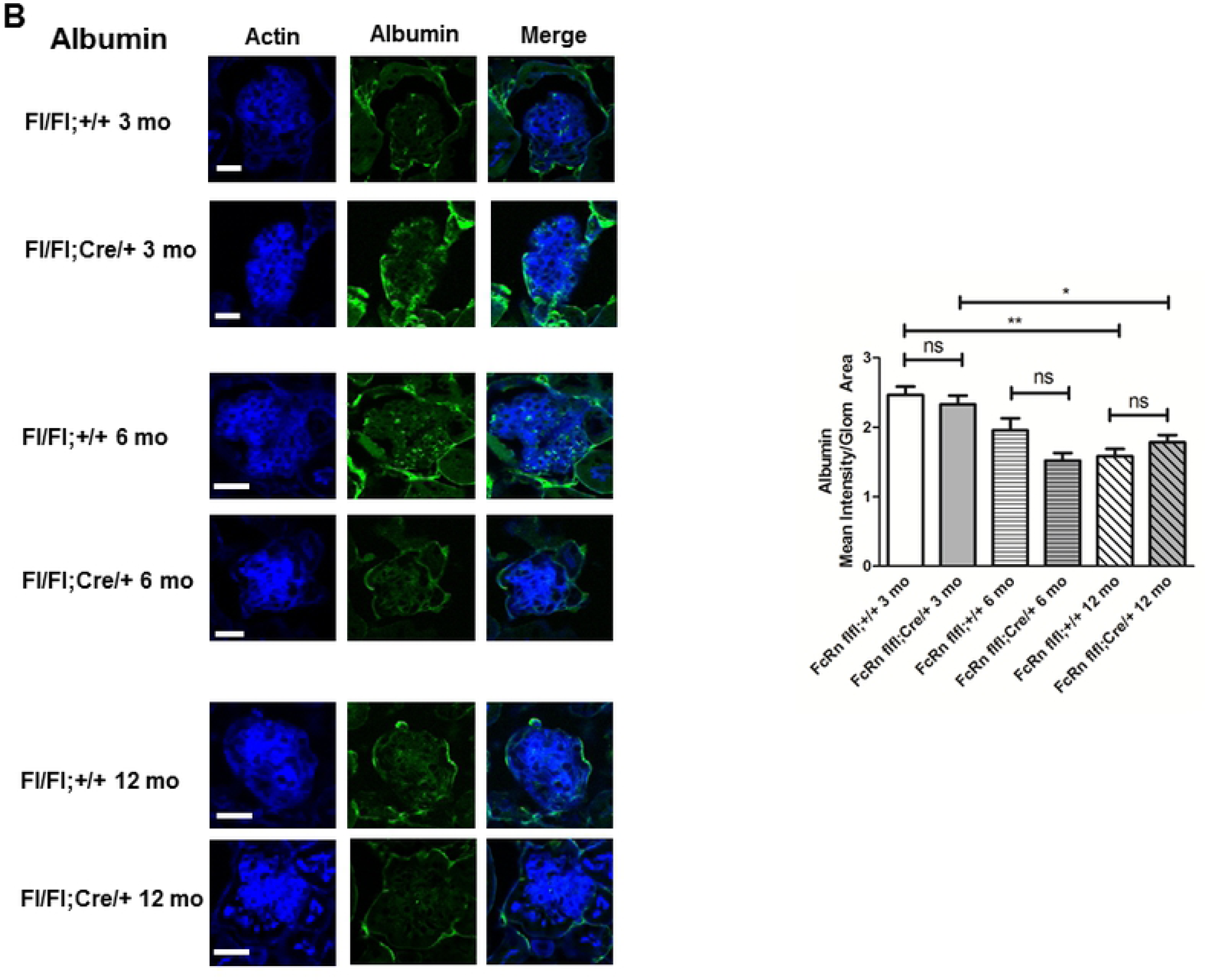
*Podocyte-specific FcRn KO resulted in a significant increase in intraglomerular IgG accumulation, but no change in albumin accumulation*. *A*, Intraglomerular IgG: At 3 months of age, there was no significant difference in IgG accumulation between podocyte specific FcRn KO (fl/fl; cre/+) and control mice (fl/fll;+/+). (n=3 mice per group). By 6 months of age there was a statistically significant increase in intraglomerular IgG accumulation in podocyte-specific FcRn KO (fl/fl;cre/+) compared to controls (fl/fl;+/+) (n= 6 control and 6 KO mice, * p < 0.05). Intraglomerular IgG accumulation was even more significantly increased by 12 months of age in the podocyte-specific FcRn KO (n= 6 control and 6 KO mice, ** p < 0.0001). Scale bar 20 μm. *B*, Intraglomerular Albumin: Albumin accumulation in control and podocyte-specific FcRn KO mouse glomeruli was minimal and was not significantly different between control and KO animals at 3, 6 or 12 months. By 12 months of age, both control and podocyte-specific FcRn KO mice had significantly less intraglomerular albumin than 3 month old control or KO animals, * p < 0.01, ** p < 0.0001. Scale bar 20 um. NS = not significant. Number of mice per group was the same as in *A*.

### Podocyte-specific KO of FcRn results in minimal intraglomerular albumin accumulation

When mean albumin intensity per glomerulus was measured, there was no significant difference in albumin accumulation in the glomeruli of 3 month, 6 month or 12 month old versus podocyte-specific FcRn KO versus control mice (Figure 4B). Interestingly, there was a significant time dependent decrease in intraglomerular albumin accumulation with 12 month old control and podocyte-specific FcRn KO mice exhibiting a significant decrease in albumin accumulation within the glomerulus compared to the respective 3 month old mice (mean albumin fluorescence/glomerular area for 12 month versus 3 month old mice 1.5 ± 0.1 vs 2.5 ± 0.1 for controls, p < 0.0001 and 1.8 ± 0.1 vs 2.3 ± 0.1 for KO, p < 0.01). The lack of albumin detection within the glomerulus was not due to an inability to stain for albumin as albumin within the blood vessels surround the glomerulus was readily detected (Figure 4B).

### Podocyte-specific KO of FcRn leads to increased mesangial:glomerular area ratio

By 6 months of age, podocyte-specific FcRn KO mice had a significant decrease in glomerular area compared to control mice (1703 ± 62.7 um^2^ 2198 ± 92.9 um^2^, p < 0.0001) and a significant increase in the mesangial to glomerular area (0.41 ± 0.01 vs 0.27 ± 0.01, p < 0.0001; Figure 5A). By 12 months of age, there was a further significant increase in the mesangial/glomerular area in the podocyte-specific FcRn KO compared to controls (0.46 ± 0.01 vs 0.35 ± 0.01, p < 0.0001), with a resultant increase in the glomerular area in the KO to close to that of control (Figure 5B). To further examine the mesangial expansion seen in the podocyte-specific FcRn KO mice, we examined the glomerular expression of α-smooth muscle actin (α-SMA), a marker of activated mesangial cells (22). We found an increase in glomerular α-SMA actin expression by 3 months in podocyte-specific FcRn KO versus control which was statistically significant by 6 and 12 month months (α-SMA intensity/glomerular area for 3 month KO vs control 3.2 ± 0.3 vs 1.2 ± 0.1; for 6 month KO vs control 7.5 ± 0.5 vs 4.0 ± 0.3, p < 0.01; for 12 month KO versus control 16.3 ± 1.0 vs 5.6 ± 0.3, p < 0.0001, Figure 6).

**Figure 5.**
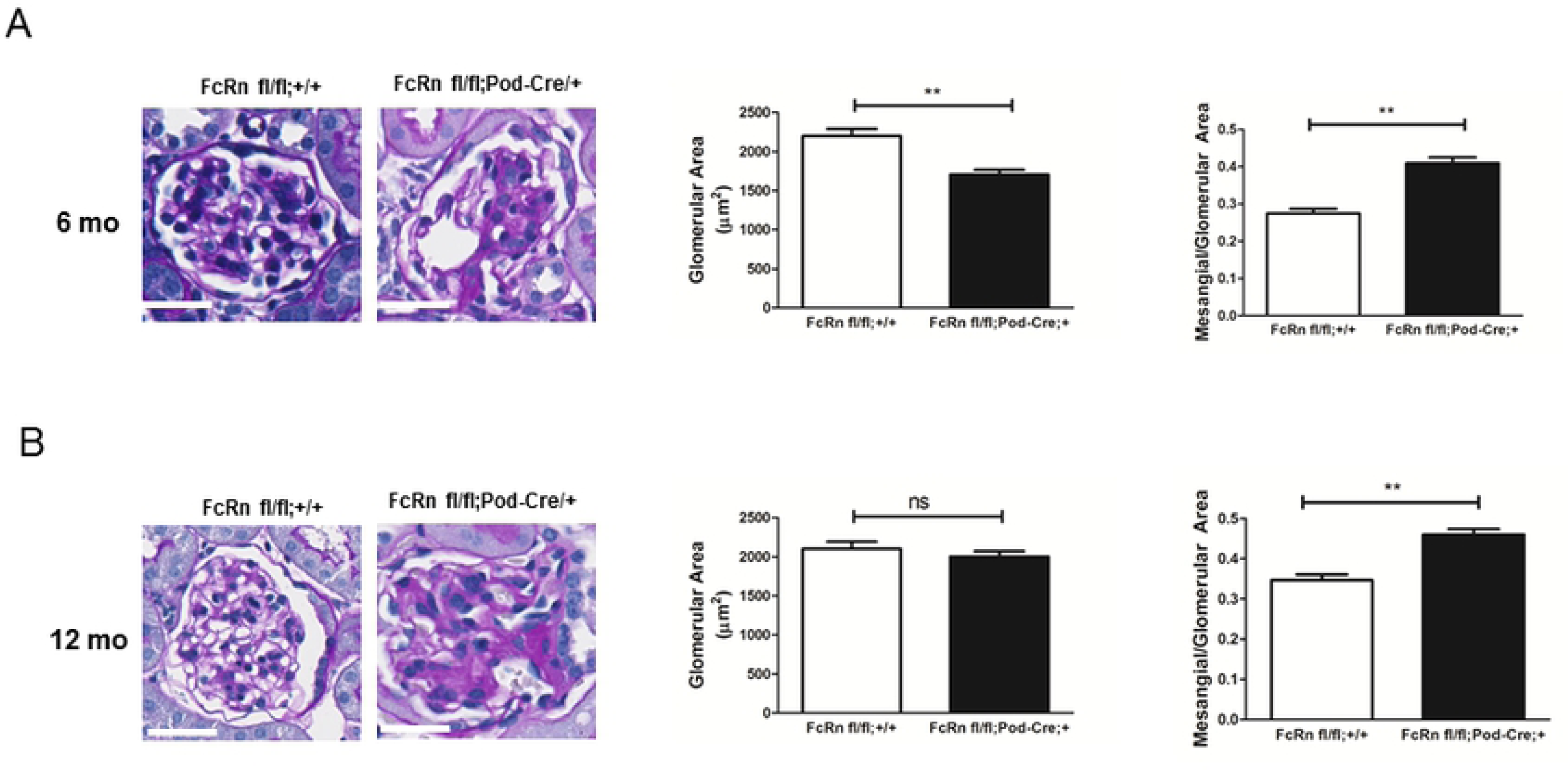
*Podocyte-specific FcRn KO resulted in mesangial expansion as mice aged*. *A*, 6 month old podocyte-specific FcRn KO mice had a decrease in mean glomerular area (**, p < 0.0001) and an increase in mesangial/glomerular area (**, p < 0.0001 compared to controls). Scale bar 20 um. *B*, By 12 months of age, mean glomerular area was similar in podocyte-specific FcRn KO and control mice but the podocyte-specific KO manifested a further increase in glomerular/mesangial area (**, p < 0.0001 compared to controls). Scale bar 20 um.

**Figure 6.**
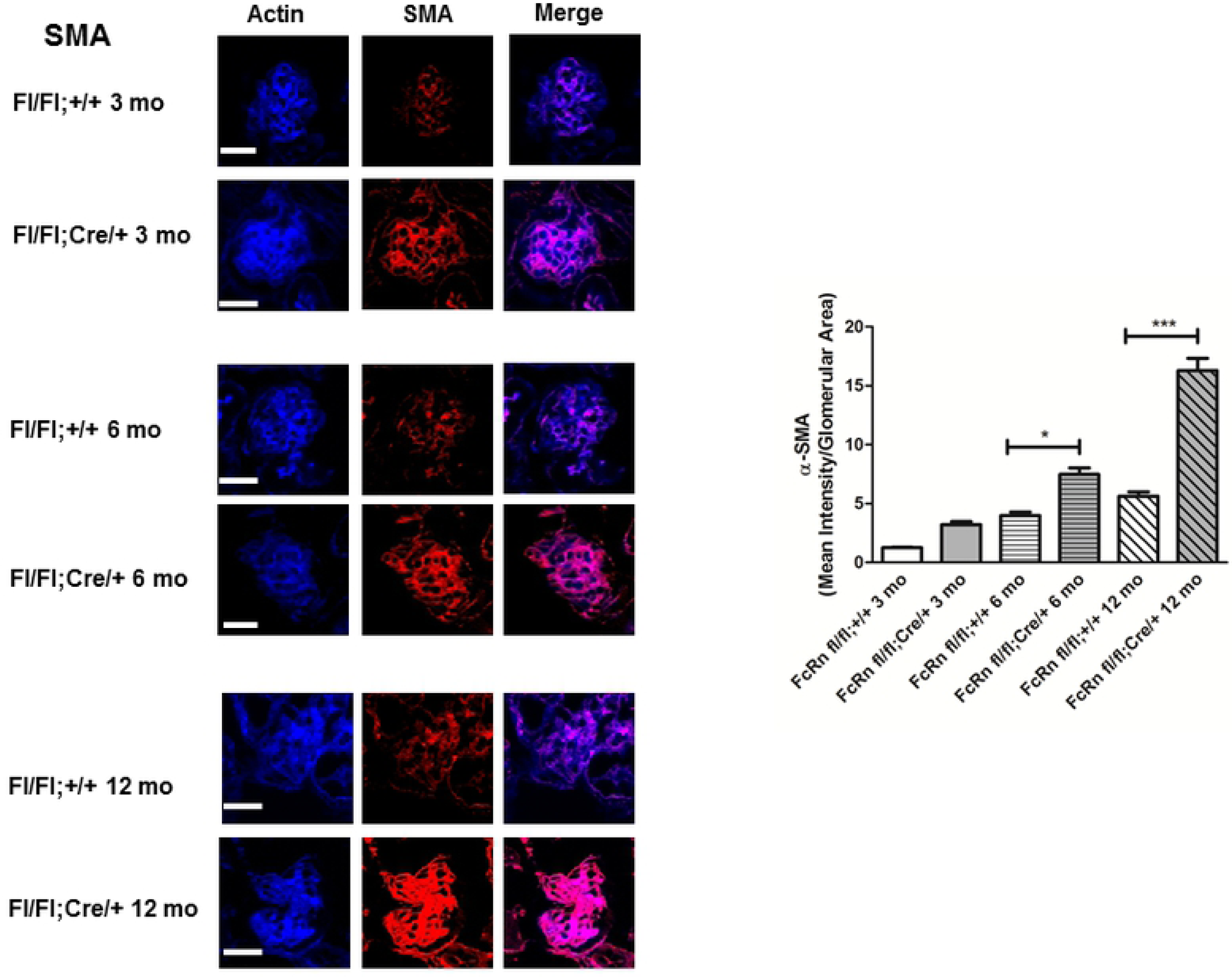
*Podocyte-specific FcRn KO resulted in increased glomerular expression of α-smooth muscle actin*. Intraglomerular expression of α-smooth muscle actin (a-SMA) was significantly increased in 6 month podocyte-specific FcRn KO (fl/fl;Cre/+) mice compared to controls (fl/fl;+/) (*, p < 0.01, n = 6 control and 6 KO mice). The increase in a-SMA expression was even more pronounced by 12 months of age (**, p < 0.0001, n = 6 control and 6 KO mice). Scale bar: 20 μm.

## Discussion

Proteinuria is a common clinical marker of kidney damage and is strongly associated with progression of kidney disease (2, 23). The mechanisms underlying renal handling of serum proteins such as albumin and IgG remain to be fully elucidated. Human and animal studies have shown podocyte vacuolization in proteinuric kidney diseases and albumin and IgG have been shown to colocalize with podocyte vacuoles (5, 10, 24-29). Previously, we have shown that cultured podocytes endocytose albumin and that the majority of endocytosed albumin is transcytosed. Here we extend our findings to an in vivo model and also examine podocyte handling of IgG. To our knowledge, this is the first systematic concurrent examination of podocyte albumin and IgG trafficking in podocytes.

In renal proximal tubular cells, FcRn is required for salvaging albumin and IgG from the degradative pathway and transcytosing these proteins from the apical cell surface to the basolateral side (15). Others have shown that injection of an anti-FcRn antibody reduces proteinuria in nephrotic rats (26) and that global FcRn KO mice accumulate IgG within the glomerulus by 6 months of age (19). Our study, however, is the first to directly demonstrate the role of FcRn in albumin and IgG handling in cell culture and in an in vivo model in which only podocytes lack FcRn. Our studies demonstrate that albumin and IgG are differentially handled by podocytes in vitro and in vivo. Lack of FcRn in cultured podocytes did not impair albumin handling acutely. In vivo, lack of FcRn did not result in significant intraglomerular accumulation of albumin. Interestingly, mice had significantly less intraglomerular albumin accumulation at 12 months compared to 3 months in both podocyte-specific FcRn KO animals and controls. Lack of intraglomerular albumin accumulation in podocyte-specific FcRn KO mice was not due to technical issues with staining as albumin could be seen in the peritubular and periglomerular capillaries. The lack of albumin accumulation in podocytes both in vivo and in vitro suggests that podocytes possess another non-FcRn dependent pathway for handling any serum albumin that passes through the GFB.

Knockout of FcRn in podocytes led acutely to intracellular accumulation of IgG in vitro and accumulation of IgG in vivo which became significant by 6 months of age. Lack of FcRn, however, did not completely abrogate IgG transcytosis in cultured podocytes as evidenced by the fact that some IgG appeared in the supernatant in FcRn KO cells after loading podocytes with IgG and incubating in IgG free solution, suggesting the existence of an additional non-FcRn dependent pathway for IgG handling.

Akilesh et al. found increased IgG accumulation in the glomeruli of global FcRn KO mice compared to wild type at 6 months of age (19). There are several important differences between our study and that of Akilesh et al. We used podocyte-specific FcRn KO mice which have normal circulating levels of albumin and IgG whereas Akilesh et al. used global FcRn KO mice which have serum albumin levels that are 50% lower and serum IgG levels that are 80-90% lower than those of wild type animals. In addition, we performed systematic quantitation of albumin and IgG staining in podocyte-specific FcRn KO mice or controls as mice aged using confocal microscopy whereas Akilesh et al. used epifluorescence microscopy on wild type or global FcRn KO mice at a single time point (6 months) and did not perform any quantitation.

An interesting finding of the present study is that podocyte-specific knockout of FcRn leads to mesangial expansion in the KO mice as they age as well as increased expression of α-smooth muscle actin within the mesangial regions of the glomerulus, suggesting activation of mesangial cells with intraglomerular IgG accumulation. The mechanisms underlying how knockout of a trafficking protein in a podocyte can lead to an expansion of mesangial area and increased expression of a-SMA remain to be further investigated.

In summary we have directly examined the role of FcRn in albumin and IgG trafficking in poodcytes and found that FcRn-mediated trafficking of these proteins differs. Our findings suggest that intra-podocyte trafficking pathways are complex and that disruption of normal trafficking pathways in podocytes is deleterious.

## Methods

### Cell culture

#### Generation of conditionally immortalized WT and FcRn KO podocytes

Podocytes were isolated from wild type or global FcRn KO mice as previously described (30). Primary podocytes were immortalized using a thermosensitive SV40 T antigen as previously described (21). Briefly, media containing viral particles was collected from the viral producer line plpcx SVtsa58 (kindly provided by Dr. Parmjit Jat) and applied to primary WT or FcRn KO podocytes. The plpcs SVtsa58 viral producer line encodes the thermolabile tsA58 LT antigen and G418 resistance. Podocytes were selected using G418. After selection, podocytes were allowed to replicate at 33 °C. To induce differentiation, podocytes were placed at 37 °C for 8 -10 days. To verify expression of podocyte markers, podocytes were stained with podocin or WT1.

#### In vitro trafficking assay

The in vitro albumin and IgG trafficking experiments were performed as previously described (12). Briefly, WT or FcRn KO podocytes were loaded with 1.5 mg/ml FITC-human albumin or 1 mg/ml human IgG at 4°C (which permits binding and inhibits endocytosis) or 37°C (which permits endocytosis). Previous work has shown that mouse FcRn binds both human albumin and IgG at the concentrations used in these studies (31). After loading, cells were washed well and incubated in Ringer solution (122.5 mM NaCl, 5.4 mM KCl, 1.2 mM CaCl_2_, 0.8 mM MgCl_2_, 0.8 mM Na_2_HPO_4_, 0.2 mM NaH_2_PO_4_, 5.5 mM glucose, and 10 mM HEPES; pH 7.4) at 4 °C or 37 °C in the presence or absence of 20 uM leupeptin (which inhibits lysosomal degradation). Cells and supernatant were either harvested immediately (t = 0) or at the designated time points. The supernatant was removed by evaporation under a vacuum and the remaining albumin or IgG resuspended in 35 ul RIPA buffer. The amount of albumin or IgG in the cellular or fraction was assessed by western blot analysis.

### Western blotting

For podocytes lysed in RIPA buffer, protein concentrations were measured by BCA assay (Pierce, Thermofisher Scientific, Waltham, MA), and samples were reduced (10% β-mercaptoethanol). Cell lysates were run on 10% polyacrylamide gels and transferred onto nitrocellulose membranes (Bio-Rad, Hercules, CA). Subsequent blocking, antibody, and wash solutions were diluted in PBS-T (phosphate-buffered saline, 1% Triton-X 100). Membranes were initially blocked (5% nonfat dry milk; 60 min) and then incubated with primary antibody. Primary antibodies include FITC (1:1,000; clone ZF2471–1900, Invitrogen; Carlsbad, CA), IgG (1:1000; GW20083F, Sigma-Aldrich, St. Louis, MO) actin (1:5,000; A1978, Sigma-Aldrich), FcRn (1:100; H-247, Santa Cruz Biotech, Dallas, TX). Blots were then washed, incubated with horseradish peroxidase-conjugated secondary antibodies (1:10,000 dilution; Jackson ImmunoResearch, West Grove, PA), and washed. The antibody complexes were detected using enhanced chemiluminescence (Pierce) and captured using a photodocumentation system (UVP; Upland, CA).

### PCR

Total RNA was isolated using RNeasy Mini Kit (QIAGEN, Valencia, CA). cDNA was synthesized from total RNA (1 µg) with the High Capacity cDNA Reverse Transcriptase Kit (Invitrogen) which uses the random primer scheme. The primers used for FcRn are as follows: sense 5′-TGA CCT GTG CTG CTT TCT CCT-3′, antisense 5′-CAG CAA TGA CCA TGC GTG GAA-3′. Real-time PCR was performed with the use of the AppIied Biosystems Step One Plus Real-Time PCR System (Life Technologies, Carlsbad, CA). The expression of a target gene in relation to a reference gene was calculated using a comparative cycle threshold (Ct) method.

### Animals

Podocyte specific FcRn knockout mice were obtained by crossing FcRn floxed mice (32) (a kind gift of Dr. Sally Ward, UT Southwestern) with podocin-Cre mice (Jackson Labs, Bar Harbor, Maine). Genotype was determined by PCR. All experimental mice were homozygous for the floxed FcRn gene. Podocyte specific FcRn knockout mice (FcRn fl/fl;cre/+) were double transgenic resulting in no FcRn expression in podocytes. Control mice (FcRn fl/fl;+/+) were single transgenic (no Cre expression) resulting in unchanged FcRn expression in podocytes. Male mice were used for all experiments. All procedures involving animals were performed using protocols approved by the Institutional Animal Care and Use Committee at the University of Colorado, Denver, protocol number 00085. Animals were euthanized using pentobarbital.

Urine albumin was measured using the Albuwell assay (Exocell), urine creatinine was measured using the assay and BUN was measured on an Alpha Wasserman auto analyzer. Serum albumin was measured by ELISA (Abcam) as was serum IgG (Affymetrix).

### Immunofluorescence

Confocal microscopy images were acquired using Zeiss 780 laser-scanning confocal/multiphoton-excitation fluorescence microscope with a 34-Channel GaAsP QUASAR Detection Unit and non-descanned detectors for two-photon fluorescence (Zeiss, Thornwood, NY). The imaging settings were initially defined empirically to maximize the signal-to-noise ratio and to avoid saturation. In comparative imaging, the settings were kept constant between samples. Images were obtained with a Zeiss C-Apochromat 40x/1.2NA Korr FCS M27 water-immersion lens objective. The illumination for imaging was provided by a 30mW Argon Laser using excitation at 488 nm, HeNe 5mW (633 nm) and 1mW (543 nm). Image processing was performed using Zeiss ZEN 2012 software. Images were analyzed in Image J software (NIH, Bethesda, Maryland). Fluorescence intensity of albumin or IgG was normalized to glomerular area. 20 -25 glomeruli were analyzed per mouse.

For fixed-cell images, podocytes were fixed in 4% paraformaldehyde in phosphate-buffered saline (PBS) with 0.5% Triton X-100 (20 min; room temp), washed, blocked with 10% normal serum and labeled with primary antibodies. Primary antibodies include: Podocin (1:200, P0372, Sigma-Aldrich), WT-1 (1: 200, sc-192, Santa Cruz), FcRn (1: 100, sc-66892, Santa Cruz). Cells were subsequently washed and labeled with the appropriate conjugated secondary antibodies (Alexa Fluor 488, Alexa Fluor 568; Invitrogen). F-actin was concurrently stained with Alexa-Phalloidin 633 (Invitrogen).

For the in vivo immunolocalization studies, the kidneys were cleared of blood by perfusion of phosphate-buffered saline (PBS) and then fixed by perfusion with 4% paraformaldehyde (Electron Microscopy Sciences; Hatfield, PA) in PBS (pH 7.4). The kidneys were then removed, immersed in 4% paraformaldehyde for 24hr, infused with 5% (2 hr), 10% (2 hr) and 25% (overnight) sucrose, frozen in liquid nitrogen and cryosectioned (3 μm). Kidney sections were blocked (10% normal goat serum in PBS) and incubated overnight at 4 °C with primary antibody: IgG (1:250; GW20083F, Sigma-Aldrich), albumin (1:250; ab106582, Abcam, Cambridge, UK), a-SMA (1:250; 1A4, Sigma-Aldrich). After washing, the sections were incubated (60 min, room temperature) with appropriate mix of Alexa 488-conjugated goat anti-chicken IgG (1:500; Invitrogen) and Alexa 633-conjugated phalloidin (1:200; Invitrogen). Sections were then washed with PBS and mounted in Fluromount-G (Thermo Fisher Scientific, Waltham, MA).

### Histology

3 um sections were cut from paraffin embedded tissue and stained using the periodic acid Schiff reagent. Analysis of glomerular and mesangial area was performed using NDP.view 2 (Hamamatsu, Hamamatsu City, Japan). 20 – 30 glomeruli were analyzed per mouse.

### Data analysis

Data are presented as means ± SE. Statistical analysis was performed using *t*-tests for two groups and one-way analysis of variance for three or more groups, using Prism software (GraphPad, San Diego, CA). Tukey’s post hoc test was applied to the ANOVA data. Values were considered statistically significant when *p* < 0.05

## Acknowledgements

This work was funded by a Norman Coplon Satellite Healthcare grant and NIH R01DK104264 to JB.

## Supplemental Figure Legends

**Figure 1.**
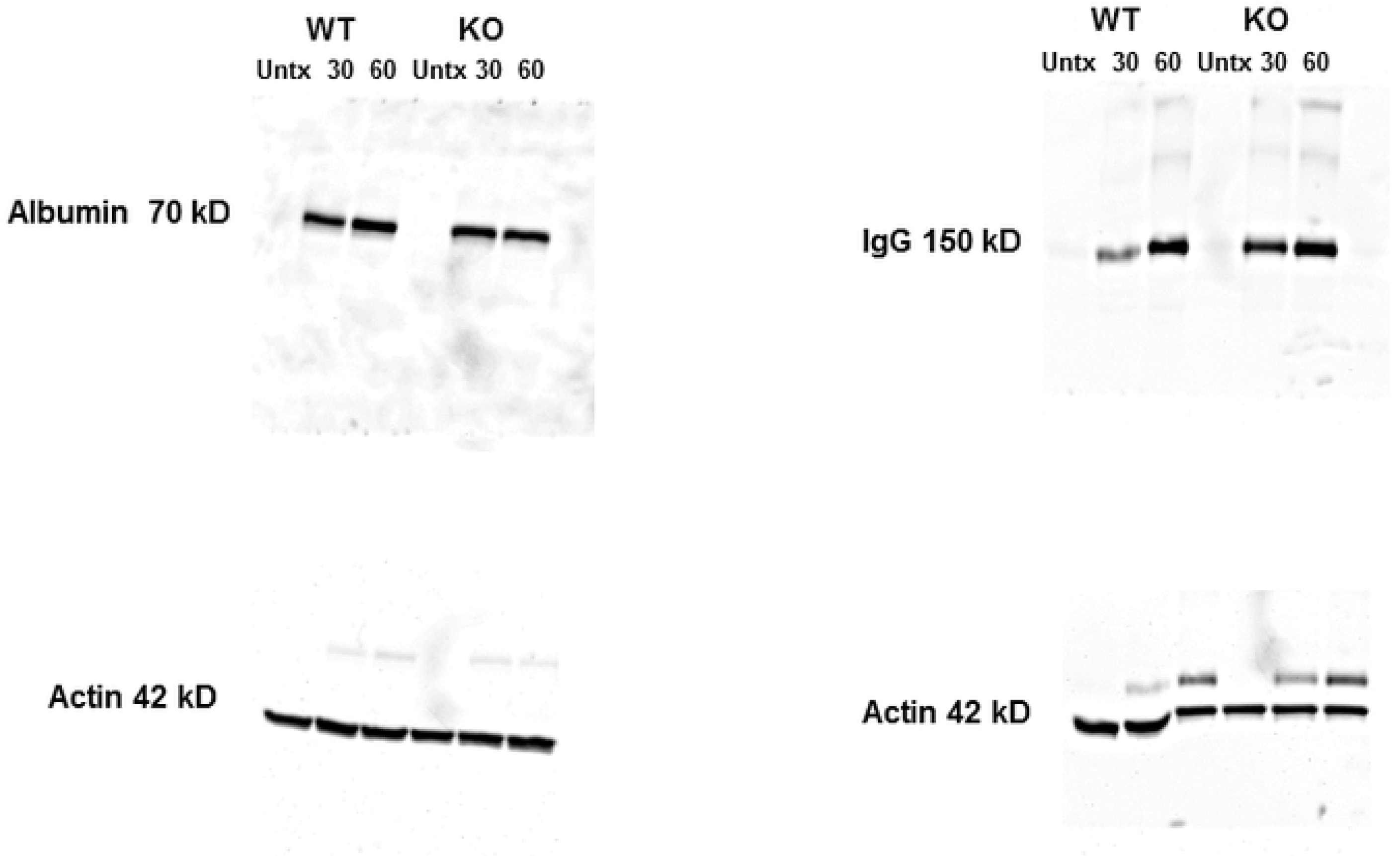
Uncropped images for western blots in Figure 1C.

**Figure 2.**
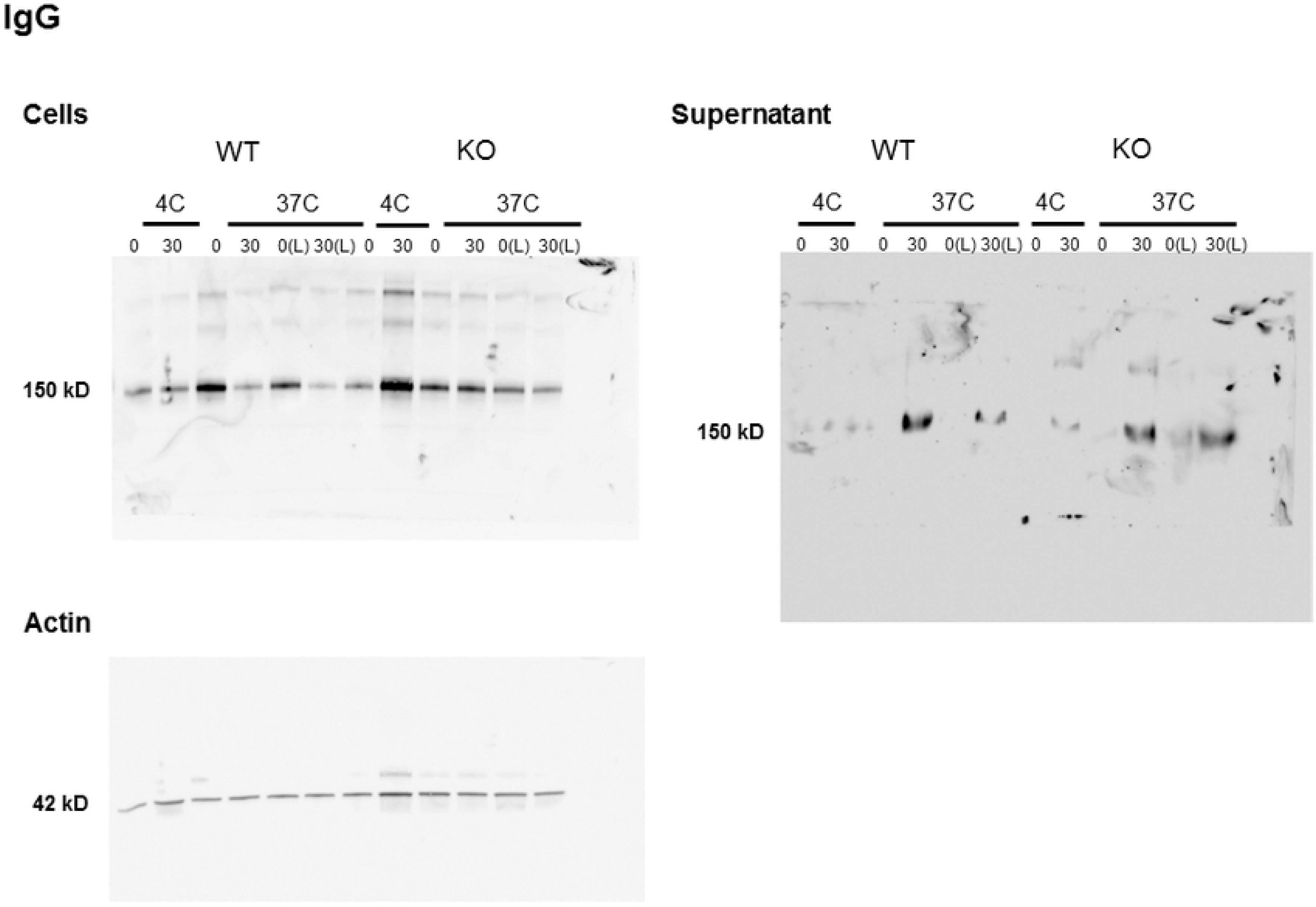
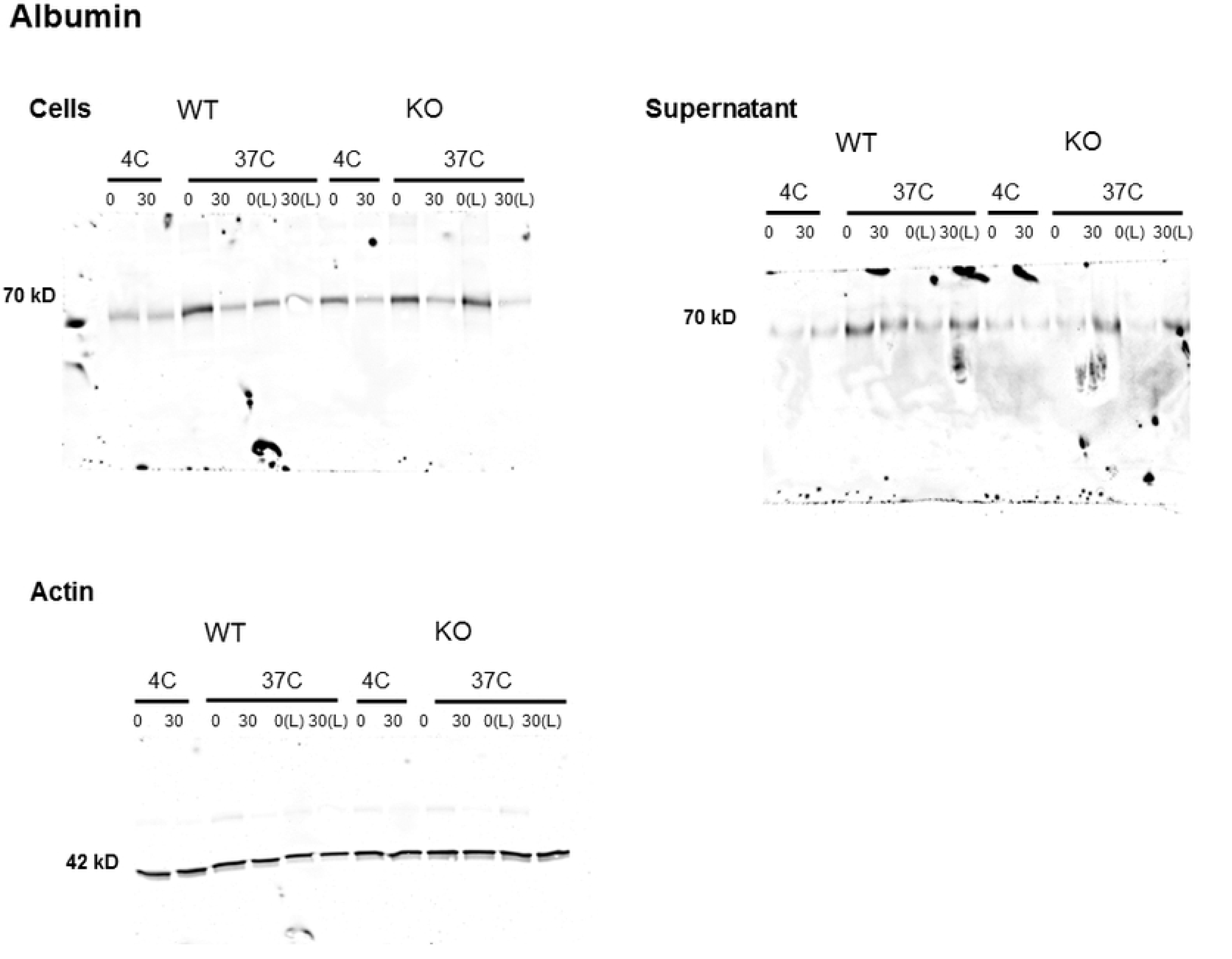
Uncropped images for western blots in Figure 2A and B.

